# The PAZ domain of *Aedes aegypti* Dicer 2 is critical for accurate and high-fidelity size determination of virus-derived small interfering RNAs

**DOI:** 10.1101/2024.06.20.599909

**Authors:** Melinda Reuter, Rhys H. Parry, Melanie McFarlane, Rommel J. Gestuveo, Rozeena Arif, Alexander A. Khromykh, Benjamin Brennan, Margus Varjak, Alfredo Castello, Lars Redecke, Esther Schnettler, Alain Kohl

## Abstract

The exogenous siRNA (exo-siRNA) pathway is a critical RNA interference response involved in controlling arbovirus replication in mosquito cells. It is initiated by the detection of viral long double-stranded RNA (dsRNA) by the RNase III enzyme Dicer 2 (Dcr2), which is processed into predominantly 21 nucleotide (nt) virus-derived small interfering RNAs, or vsiRNAs that are taken up by the Argonaute 2 (Ago2) protein to target viral single-stranded RNAs. The detailed understanding of Dicer structure, function and domains owes much to studies outside the context of viral infection, and how Dcr2 domains contribute to detecting viral dsRNA to mount antiviral responses in infected mosquito cells remains much less understood. Here, we used a Dcr2 reconstitution system in *Aedes aegypti* derived Dcr2 KO cells to assess the contribution of the PAZ domain to induction of the exo-siRNA pathway following infection with Semliki Forest virus (SFV; *Togaviridae*, *Alphavirus*). Amino acids critical for PAZ activity were identified, and loss of PAZ function affected the production of 21 nt vsiRNAs -though not the overall ability of Dcr2 to process viral dsRNA- and silencing activity. This study establishes the importance of correct vsiRNA size in mosquito exo-siRNA antiviral responses, as well as the PAZ domain’s functional contribution to Dcr2 processing of viral dsRNA to 21 nt vsiRNAs.

## INTRODUCTION

The emergence and re-emergence of arboviruses pose a continuing threat to human and animal health, with mosquito-transmitted pathogens such as dengue virus (DENV; *Flaviviridae, Flavivirus*), Zika virus (ZIKV; *Flaviviridae, Flavivirus*) and chikungunya virus (CHIKV; *Togaviridae, Alphavirus*) as prominent examples of the threats posed (Zaid et al. 2021; Halstead 2019; Weaver et al. 2018; Weaver and Reisen 2010; Huang et al. 2019; Pierson and Diamond 2020). The mosquito *Aedes aegypti* is a critical vector for the transmission of many human-infecting arboviruses, including DENV, ZIKV and CHIKV. It is found across many areas of the tropics and subtropics where it is well adapted to the human environment, preferentially taking bloodmeals from human hosts which can result in arbovirus transmission. Prevention of transmission frequently relies on chemical control measures that target vectors, though novel approaches such as *Wolbachia* endosymbionts and gene drive-based approaches are promising (Achee et al. 2019; Facchinelli et al. 2023; Carvalho and Moreira 2017; Brady and Hay 2020; Caragata et al. 2021; Wang et al. 2021, 2022).

Arboviruses actively replicate in their vectors, which results in the induction of immune responses. Understanding these is relevant for a better comprehension of vector processes that impact arbovirus transmission and thus targets for intervention (Alonso-Palomares et al. 2018; Lambrechts and Saleh 2019). Among host responses that control arbovirus replication in vectors, the RNA interference response -specifically the exogenous siRNA (exo-siRNA) pathway-plays a critical role. Current understanding of this pathway comes from early studies in *Drosophila melanogaster*, with critical components and mechanisms also conserved in mosquitoes thus, proving the importance of the exo-siRNA pathway across insects. It is initiated following targeting of viral replication induced long double-stranded RNA (dsRNA) by the RNase III enzyme Dicer 2 (Dcr2), which cleaves the dsRNA into virus-derived small interfering RNAs (vsiRNAs). In mosquitoes and *D. melanogaster*, each vsiRNA strand is predominantly 21 nucleotides (nt) in length, with a 19 nt overlap and 2 nt overhangs at the 3’end. In the next step, 21 nt vsiRNAs are taken up by Argonaute 2 (Ago2). Ago2 is part of the RNA-induced silencing complex (RISC) and retains one strand of the siRNA to target and degrade complementary viral RNAs (Blair and Olson 2015; Tikhe and Dimopoulos 2021; Olson and Blair 2015; Bronkhorst and van Rij 2014; Swevers et al. 2018; Prince et al. 2023; Leggewie and Schnettler 2018; Samuel et al. 2018). Indeed, absence of Dcr2 in mosquito cell lines resulted in the loss of 21 nt vsiRNAs and impaired antiviral responses (Brackney et al. 2010; Gestuveo et al. 2022; Scott et al. 2010; Varjak et al. 2017), while *Ae. aegypti* mosquitoes without functional Dcr2 develop disease phenotypes, increased replication and dissemination following arbovirus infection (Merkling et al. 2023; Samuel et al. 2023). These data show the importance of Dcr2 as an initiator of the exo-siRNA pathway, but other functional aspects of this effector protein are less understood. The domain structure of *Ae. aegypti* Dcr2 (Aaeg Dcr2) is similar to that of *D. melanogaster* Dcr (Dm Dcr2), with helicase-DUF-PAZ-RNase 3A-RNase 3B-dsRBD from N to C terminus of the protein (Fig. 1), though other functional domains are present in the protein, too (Paturi and Deshmukh 2021). Structural studies have revealed that Dicer proteins generally have an L shape in which these different domains are arranged (Paturi and Deshmukh 2021; Zapletal et al. 2023; Ciechanowska et al. 2021). Analyses of Dm Dcr2 have confirmed this shape (Yamaguchi et al. 2022; Sinha et al. 2018; Su et al. 2022), which was also predicted for Aaeg Dcr2 using AlphaFold (Gestuveo et al. 2022). The bottom (or base) module of the L-shaped Dcr2 structure contains the helicase domain; the core comprises RNase 3A/B and dsRBD domains, and the cap module the PAZ (Piwi/Ago/Zwille) domain. Biochemical and structural studies have investigated the dsRNA binding properties of Dm Dcr2 domains, with the PAZ domain preferably binding dsRNA termini with 3’ overhangs (Sinha et al. 2018). The structural arrangements of the 3’ nt overhang binding pocket in the PAZ domain and 5’ monophosphate in the adjacent Platform domain have been investigated in detail.

**Figure 1.**
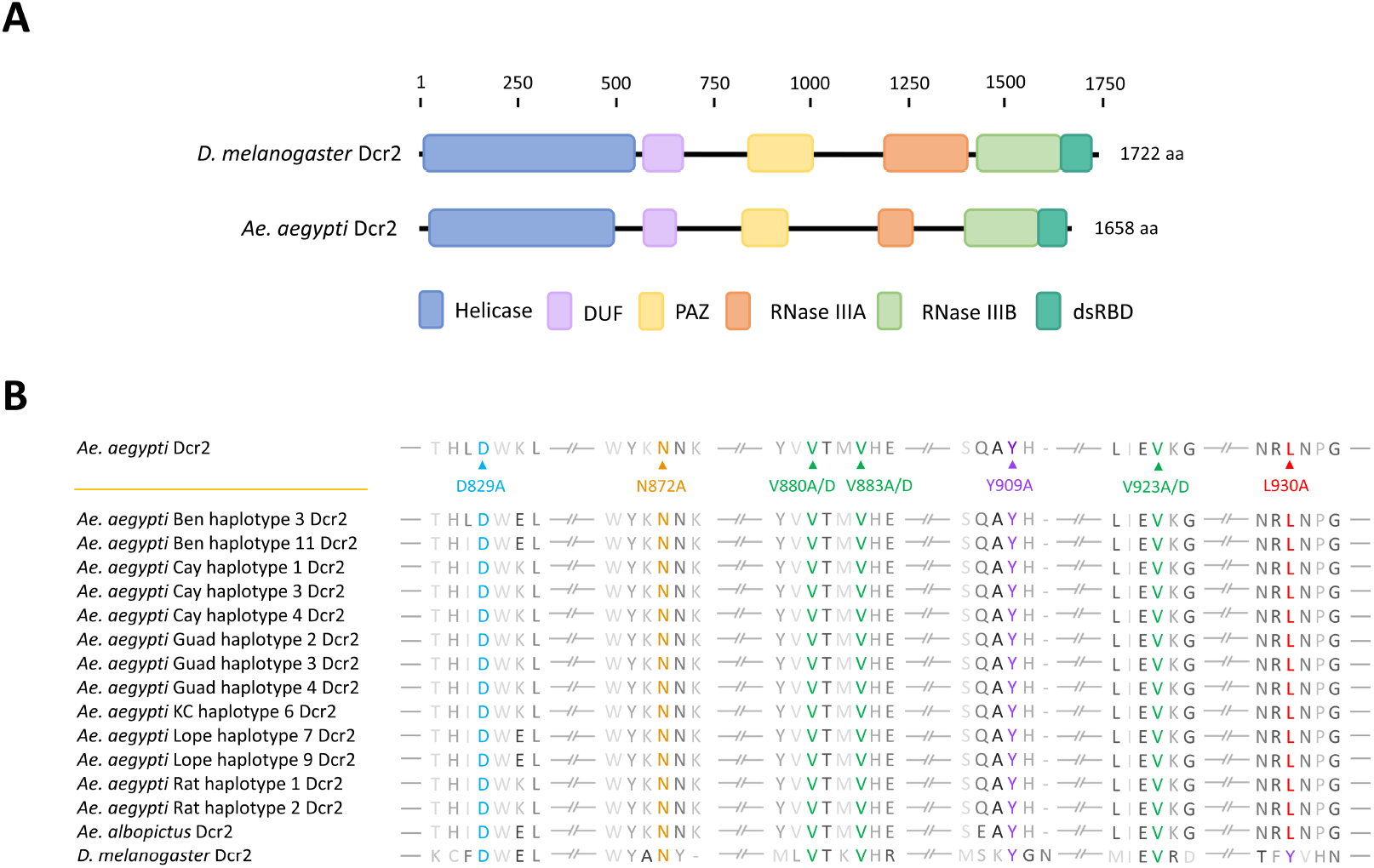
Domain features of Dcr2. **(A)** Schematic representation of *D. melanogaster* and *Ae. aegypti* Dcr2, including functional domains: Helicase domain, domain of unknown function (DUF), PIWI-Argonaute-Zwille (PAZ) domain, RNase IIIA and IIIB domains, and dsRNA binding domain (dsRBD); aa, amino acid **(B)** Multiple sequence alignment of insect Dcr2 referring to Aaeg Dcr2 AAW48725, including the selected mutations of highly conserved amino acids for potential PAZ domain loss of function. Accession numbers for sequences used are indicated in Supplemental Table S1. Alignment was produced using Benchling (https://www.benchling.com/).

Importantly, interactions of dsRNA with PAZ and Platform via 3’ 2 nt overhang and 5’ monophosphate of the dsRNA allows positioning and spacing of the dsRNA for precise dicing activity and siRNA length (Su et al. 2022; Yamaguchi et al. 2022). These structural data consolidate a previous study on the importance of binding of the 5’ monophosphate in dsRNA for length fidelity in 21 nt siRNA production from long dsRNA-though not efficiency of small RNA production *per se*; and mutagenesis analysis of the Dm Dcr2 PAZ domain also demonstrated the critical role of 21 nt siRNAs in silencing an inverted repeat transgene in the fly model (Kandasamy and Fukunaga 2016). Interestingly, the presence of 5’ monophosphate and two specific arginine residues in Dm Dcr2 PAZ were found to be critical for cleavage of short dsRNA (Cenik et al. 2011; Fukunaga et al. 2014), though not longer ones where (as stated above) accurate length determination of siRNAs is the critical contribution of 5’ monophosphate binding to the dicing process.

Studies on the insect Dcr2 PAZ domain relied on *in vitro* studies with dsRNA substrates or endogenous small RNAs, while the role in antiviral responses during infection, to our knowledge, is still unclear. The importance of the Dcr2 helicase domain in antiviral responses has been previously shown (Gestuveo et al. 2022; Marques et al. 2013; Donelick et al. 2020), there remains a critical gap because the nature of the viral dsRNA substrate is unknown-does Dcr2 detect long, short, or different types of dsRNA in infected cells, and how does recognition occur? Additionally, the most efficient siRNA size in mosquito cells is only assumed to be 21 nt but not definitively shown. Here we aimed to investigate the role of the Aaeg Dcr2 PAZ domain in mosquito antiviral vsiRNA generation and its requirement for efficient antiviral responses. For this, we used the previously described Aaeg Dcr2 activity reconstitution system in *Ae. aegypti* derived Dcr2 KO cells, with which we demonstrated the importance of the helicase and RNase domains and that 21 nt vsiRNAs are generally produced by a processive production mechanism from viral dsRNA (Gestuveo et al. 2022). To understand the contribution of the PAZ domain to the antiviral activity of Aaeg Dcr2, we identified conserved amino acids in Aaeg Dcr2 PAZ which were mutated to assess their impact on dsRNA processing activity. Mutations abrogated antiviral activity against a positive sense RNA virus, Semliki Forest virus (SFV), a member of the *Togaviridae* family, genus *Alphavirus*. PAZ function inactivating mutations resulted in a loss of 21 nt vsiRNA generation, but not a loss of the small RNAs of other sizes. These findings indicate that PAZ domain functionality is critical for determination of the usually predominant vsiRNA length, with 21 nt optimal for antiviral responses, but mutations did not affect processive cleavage of viral dsRNA as such.

## RESULTS

### Identification of amino acids in Aaeg Dcr2 PAZ domain required for antiviral activity

Analysis of Aaeg Dcr2 sequence (InterProScan, Geneious) indicated that the PAZ domain was located from positions N824 to G941 (Fig. 1A). This was in line with NCBI domain analysis, which also located the PAZ domain between amino acids 824-941. Next, Dcr2 sequences of *Ae. aegypti* (including recently available sequences, see (Gestuveo et al. 2022)), *Ae. albopictus* and *D. melanogaster* PAZ were aligned to identify highly conserved amino acids with potential functional activities. As shown in Fig. 1B, amino acids D829, N872, V880, V883, Y909, V923, and L930 were maintained in Aaeg Dcr2 and conserved in Dm Dcr2 as well as *Ae. albopictus* Dcr2. For completeness, corresponding amino acid positions in Dm Dcr2 are indicated in Fig. 2A. Using NCBI domain analysis, the PAZ domain of Dm Dcr2 (1722 amino acids in length) was shown as located from amino acids 843-1004.

**Figure 2.**
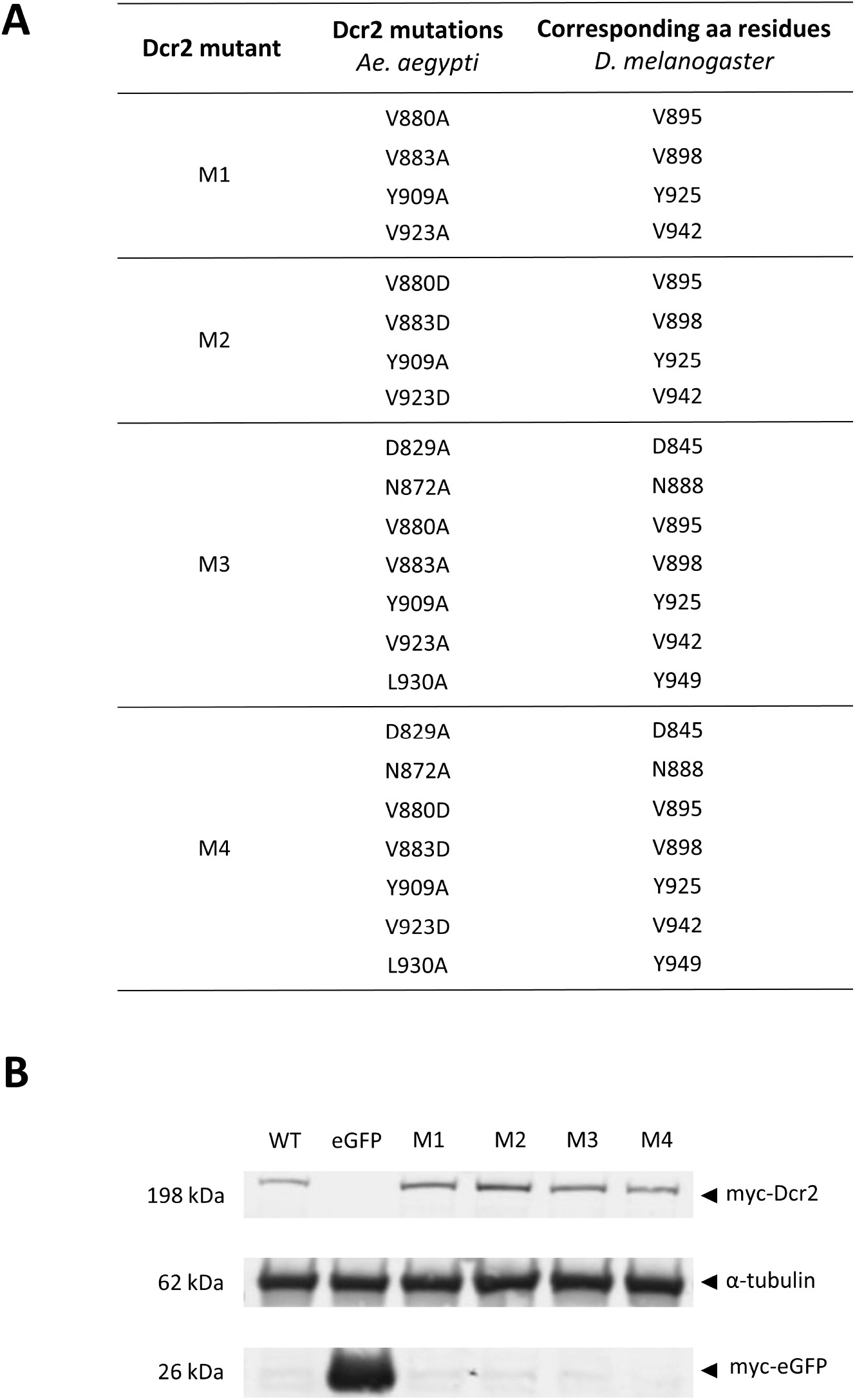
Mutation and expression of Aaeg Dcr2. **(A)** Table showing produced Aaeg Dcr2 PAZ domain mutants, the respective introduced loss-of-function mutations, as well as corresponding amino acid residues in Dm Dcr2; aa, amino acid. **(B)** Assessment of (myc-tagged) Aaeg Dcr2 expression by western blot analysis. AF319 cells were transfected with either pPUb plasmids expressing WT or PAZ domain mutant M1-M4 Dcr2, using pPUb-myc-eGFP as control. Anti-myc and anti-α tubulin (control) antibodies were used. Representative of three independent repeats (other repeats in Supplemental Fig. S2).

Alphafold was used to determine the location/structure of these conserved amino acids within the structure of the Aaeg Dcr2 PAZ domain, which appeared mainly located (with the exception of L930) around its core (Supplemental Fig. S1) when compared to Dm Dcr2 (Yamaguchi et al. 2022), suggesting structural and/ or functional importance.

Combinatorial mutagenesis was carried out as described for mutants M1-4 (conserved amino acids were replaced as indicated) (Fig. 2A). PAZ domain mutant M1-M4 and WT Dcr2 expression in AF319 Dcr2 KO cells transfected with the corresponding expression constructs was assessed by western blot, showing that all proteins were expressed (Fig. 2B, Supplemental Fig. S2).

Next, we assessed the functionality of the Aaeg Dcr2 mutants for their silencing ability and antiviral activity. WT Dcr2 displayed antiviral activity against SFV-FFLuc in this transfection-based assay (Fig. 3), as previously demonstrated (Gestuveo et al. 2022). M1-M4 Dcr2 mutants lost SFV-FFLuc antiviral activity, compared to WT Dcr2. Similarly, the functional silencing abilities of the mutant Dcr2 were investigated in a reporter plasmid-based assay, where exogenous dsRNA is provided to induce the exo-siRNA pathway against a reporter gene (Gestuveo et al. 2022). Here, M1-M4 Dcr2 mutants only retained residual (although significant compared to control) ability to silence the reporter gene. This indicated that the mutation combinations in M1-M4 all affected Dcr2 PAZ domain functionality, and thus Aaeg Dcr2’s ability as an effector in the exo-siRNA pathway.

**Figure 3.**
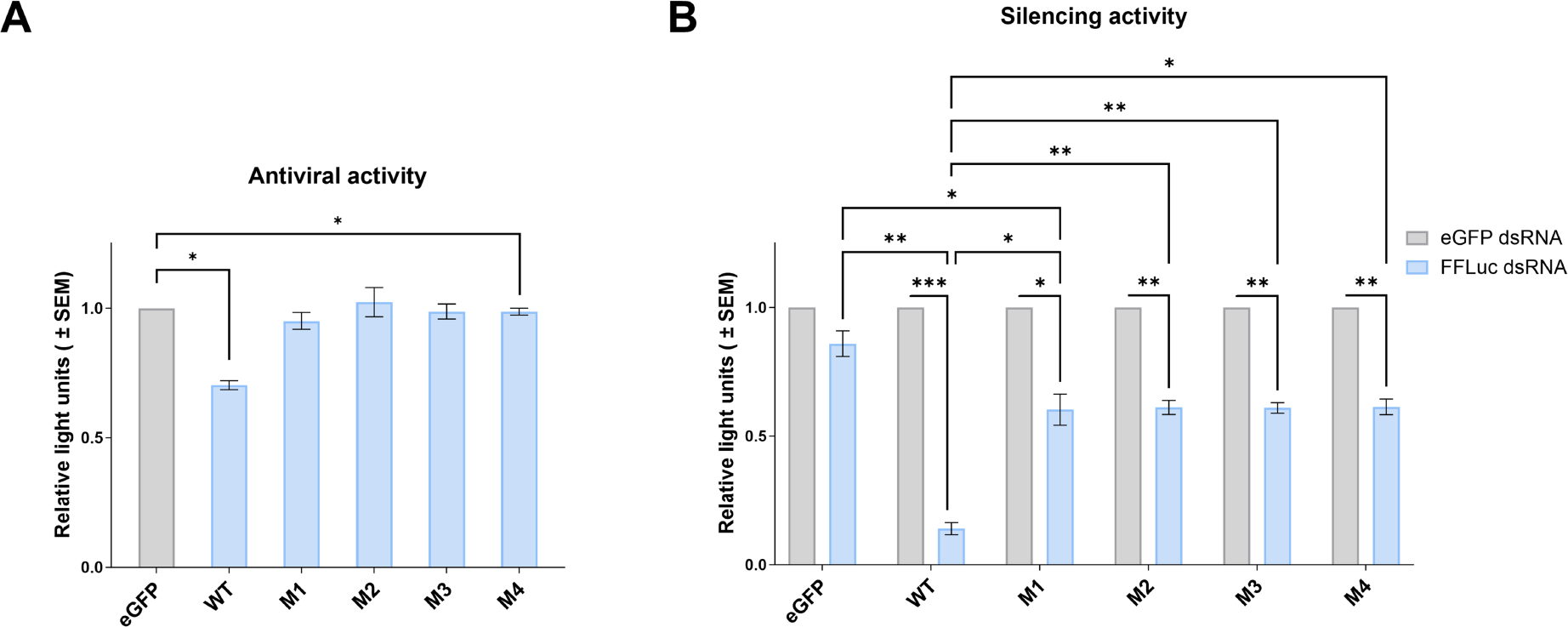
Functional analysis of Aaeg Dcr2 PAZ domain mutants. **(A)** Antiviral activity of Aaeg Dcr2. AF319 cells were transfected with expression plasmids encoding myc-tagged WT or PAZ domain mutant M1-M4 Dcr2 or eGFP (negative control). These cells were infected with SFV-FFLuc (MOI = 0.1) 24 hours post-transfection (hpt). FFLuc-levels were measured at 48 hours post-infection (hpi) and are shown as mean ± SEM relative light units compared to eGFP control set to 1 from 3 independent repeats, performed in technical triplicate with * indicating p<0.05 according to Student’s t-test. **(B)** Silencing activity of mutant Dcr2. AF319 cells were co-transfected with (i) myc-tagged pPUb plasmids expressing WT or PAZ domain mutants M1-M4 Aaeg Dcr2 (with eGFP as control), (ii) FFLuc and RLuc (internal control) reporter plasmids and (iii) dsRNA targeting FFLuc (FFLuc dsRNA, blue) or eGFP (eGFP dsRNA; non-silencing control, grey). At 24 hpt, FFLuc and RLuc levels were measured and are shown as mean ± SEM relative light units (FFLuc/RLuc) normalized to dseGFP from 3 independent repeats, performed in technical triplicate, with * = p<0.05, ** = p<0.01, *** = p<0.001 versus controls according to two-way ANOVA.

### Aaeg Dcr2 PAZ domain loss of function mutations: effects on 21 nt vsiRNA production

The impact of Aaeg Dcr2 PAZ domain mutations on vsiRNA production was investigated utilising high throughput small RNA sequencing from AF319 cells transiently expressing PAZ domain mutant or WT Dcr2 and infected with SFV. Raw populations of small RNA read lengths prototypical of miRNAs (22 nt), siRNAs (21 nt) and piRNAs (24-30 nt) (Supplemental Fig. S3) were observed in all cells. Small RNA profiles from M1-4 Dcr2 mutants and the eGFP control demonstrated a bias towards a read length of 22 nt rather than 21 nt observed in the WT Dcr2 expressing cells. Read mapping profiles of SFV-derived small RNAs indicated that WT Dcr2 efficiently mediated the production of SFV-derived 21 vsiRNAs (Fig 4), which are (as expected, see (Gestuveo et al. 2022)) the predominant length in the 18-22 nt size range and aligned along the length of the SFV genome and antigenome. Almost no such vsiRNAs were detected in the eGFP control. Importantly, M1-M4 Dcr2 mutants showed reduced production of 21 nt vsiRNAs (though those remaining align along the length of SFV4 genome and antigenome) and a more even distribution of read numbers across the 20-22 nt small RNA size range, compared to WT Dcr2, was observed. This indicates that M1-M4 Dcr2 PAZ domain mutants have lost the ability to precisely measure siRNA length and consequently do not predominantly produce 21 nt vsiRNAs; but have retained the ability to process viral dsRNA into small RNAs. Critically, AlphaFold 3 based modelling of Aaeg Dcr2 PAZ domain interactions with dsRNA suggested that mutations did not abrogate the ability to bind dsRNA (Supplemental Fig. S4). This consolidates the idea that conserved amino acid are required for keeping dsRNA in a precise position for accurate cleavage to occur.

**Figure 4.**
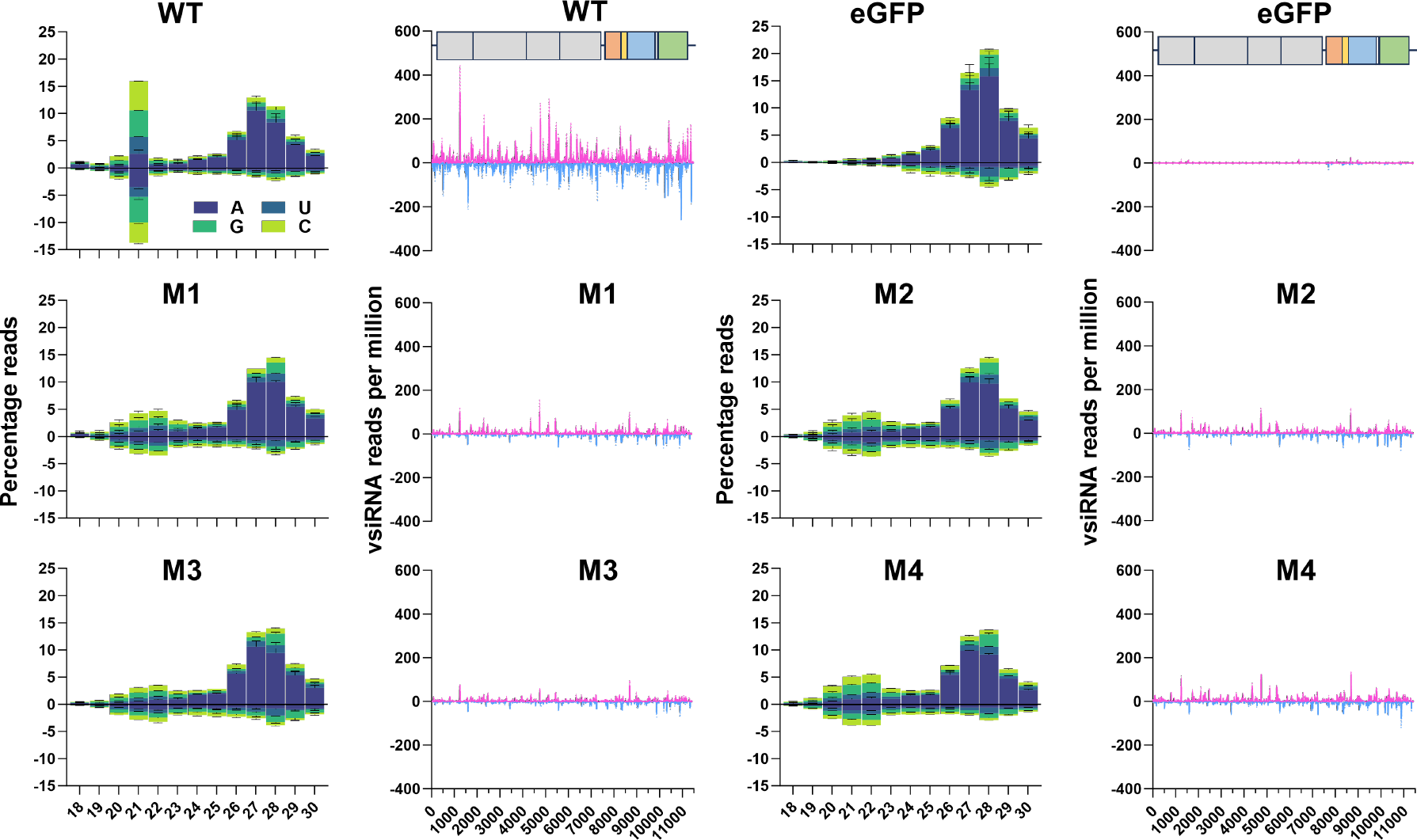
Aaeg Dcr2 PAZ domain mutations affected 21 nt vsiRNA magnitude and length. Small RNA sequencing of AF319 cells, transiently expressing PAZ domain mutant M1-M4 or WT Dcr2, or eGFP (as negative control), and infected with SFV (MOI=1) at 48 hpi. Histogram of small RNA reads 18-30 nt in length, mapped to the SFV (SFV4 GenBank ID: KP699763) genome (positive numbers) and antigenome (negative numbers) with colours indicating first base nucleotide prevalence per size, shown as mean % mapped reads (Y axis, percentage reads) from two independent experiments +/- SD. Mapping of SFV-derived 21 nt vsiRNAs, mapped along the SFV genome (magenta) or anti-genome (cyan) (Y axis, vsiRNA reads per million) +/- SD.

### Virus-derived small RNAs retain a 2 nt overhang cleavage pattern independently of Aaeg Dcr2 PAZ domain functionality

The predominant 21 nt vsiRNAs produced by Dcr2 cleavage that are commonly observed following arbovirus infection of mosquito cells are generated as duplexes. Assuming that mechanisms observed for *D. melanogaster* also apply in mosquitoes, they consist of two 21 nt strands that overlap by 19 nt with a 2 nt overhang at each 3’end. To investigate if and how dsRNA cleavage patterns were affected by the PAZ domain for generating virus-derived small RNAs, we analysed strand overlapping sRNA patterns (Fig. 5A). For WT Dcr2, but also M1-M4 Dcr2 mutants, all virus derived small RNAs in the range of 18-23 nt all showed a clear pattern, with strand complementarity significantly enhanced for overlaps that result in 2 nt overhangs. For example, 21 nt vsiRNAs overlapped with complementary sequences predominantly by 19 nt (thus confirming observations in *D. melanogaster*); virus-derived small RNAs that are 20 nt in length overlapped with sequences that are 18 nt in length etc. This showed that PAZ mutations affected Aaeg Dcr2’s ability to produce the usually predominant 21 vsiRNA over other lengths but did not affect the cleavage mechanism itself. Production of virus-derived PIWI-interacting small RNAs (vpiRNAs), which are generally longer (24-30 nt) with a 10 nt overlap of the sense and antisense piRNAs as a result of the so-called ping-pong amplification mechanism in a Dcr2 independent manner (Varjak et al. 2018), was detected across all conditions.

**Figure 5.**
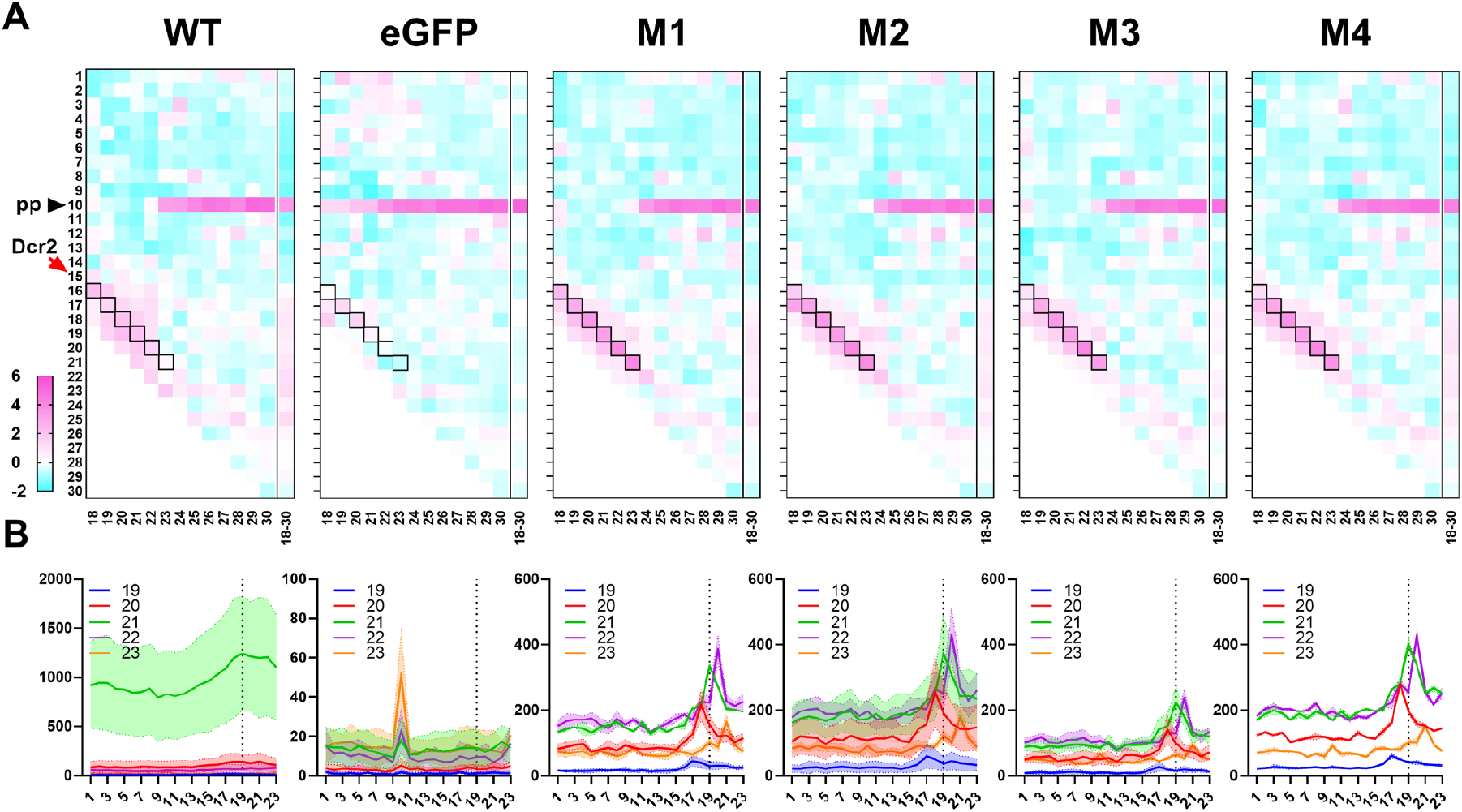
Aaeg Dcr2 PAZ domain mutations result in dysregulated vsiRNA duplex overlap lengths. **(A)** Heat maps showing mean overlap probability z-scores of 18–30 nt SFV-derived small RNAs from AF319 transiently expressing PAZ domain mutant M1-M4 or WT Dcr2, or eGFP (control) from 2 independent repeats, as initially characterized in Fig 4. Differing lengths of vsiRNAs, analysed individually, are shown horizontally, and nt overlaps are listed vertically. The red arrow labelled Dcr2 indicates the expected 2 nt overlap from dsRNA cleavage with cells boxed in black. The black arrow labeled pp (ping-pong) shows the expected 10 nt overlap from potential ping-pong amplification in the piRNA pathway. **(B)** Number of overlapping pairs per million mapped reads of 19-23 nt virus-derived small RNAs from AF319 transiently expressing PAZ domain mutant M1-M4 or WT Dcr2, or eGFP (control) infected with SFV. Data are shown as the mean of 2 independent repeats with the range of values.

### Loss of Aaeg Dcr2 PAZ domain functionality changes the position and magnitude of vsiRNA cleavage

We have previously shown that Aaeg Dcr2 produces a diverse but largely consistent pool of 21 nt vsiRNAs following SFV infection; a characteristic lost in helicase domain mutants (Gestuveo et al. 2022). To assess the impact of PAZ domain mutations on virus-derived small RNA population, diversity and cleavage bias, we examined these through two different analyses. First we defined a position-based “harmony” metric which incorporates the strand bias and magnitude of the vsiRNAs and second, using a vsiRNA population approach where each unique siRNA was treated as an individual gene and examined differential vsiRNA expression between treatment groups. While visualization of virus-derived small RNA coverage, as demonstrated in Fig. 4, provides a reasonable indicator of the origin and magnitude of vsiRNAs, a more nuanced analysis of the consistency of Dcr2 cleavage over the SFV-derived dsRNA requires consideration of the position, magnitude and strandedness of 21 nt vsiRNAs. To develop a unified metric that encapsulates these attributes at each genomic position, we calculated the terminal 5’ vsiRNA read depth at each position of the SFV genome and antigenome, and normalized the total depth to one. Subsequently, we computed the normalized differential coverage, *d_i_*, as the difference between the normalised genome (positive-sense) and anti-genome (negative-sense) coverage proportions at each position. The genome coverage proportion at position *i*, *G_i_*, is defined as the positive-sense coverage at *i* divided by the total positive-sense coverage for all *N* positions. The anti-genome coverage proportion, *A_i_*, is similarly calculated as the negative-sense coverage at *i* normalized by the total negative-sense coverage. The formula for the normalized differential coverage at each position *i* is given by:

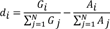

Through this measure, we obtained a ratio reflecting the balance of genome to antigenome virus-derived small RNA coverage, which provides insight into the strand-specific cleavage patterns of WT Dcr2 across the positions of the SFV genome. Using this metric, we generated a series of scatter plots (Fig. 6A) to compare the *d_i_* values for each position of the SFV genome between WT Dcr2, negative control eGFP and the PAZ domain Dcr2 mutants M1-M4. We utilized linear regression analysis to assess the relationship between *d_i_* values for all positions on the SFV genome from the WT Dcr2 and each mutant, with the coefficient of determination (R^2^) reflecting the extent to which both treatments shared related vsiRNA profile “harmony”. The regression analysis revealed varying degrees of correlation, with the lowest correlation between the eGFP control treatments compared to the WT Dcr2 (R^2^=0.115). This suggests very little positional cleavage harmony between the WT Dcr2 and eGFP control, which is to be expected. Mutants M1-M4 showed a more consistent pattern of Dcr2 cleavage to WT with R^2^ values between 0.39-0.41. Finally, we compared *d_i_* values for all 21 nt vsiRNA reads for all treatments and between mutants and summarised the R^2^ values as a heat map (Figure 6B). The results indicated that the PAZ domain mutants M1-M4 were more closely related to each other than WT Dcr2 or eGFP control with R^2^ between 0.74-0.84, indicating that the introduced mutations resulted in more harmonious cleavage patterns across the SFV genome but different from wild type.

**Figure 6.**
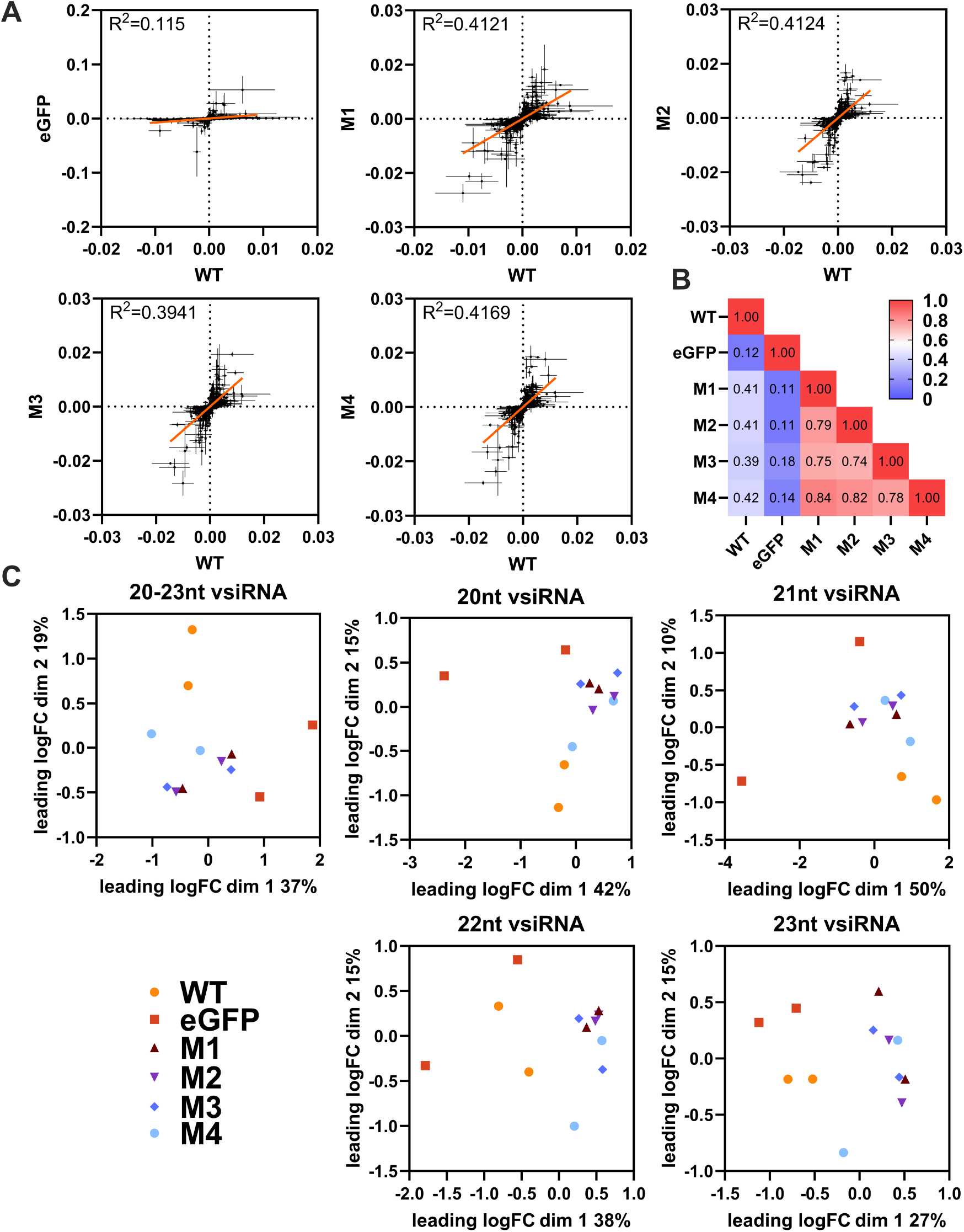
Mutations in the PAZ domain of Aaeg Dcr2 change the position and magnitude of cleavage over the SFV genome and virus-derived small RNA populations compared to WT Dcr2. Samples from Fig 4 were analysed as follows. **(A)** Scatter plots showing normalized differential coverage (*d_i_*) for each position in the SFV genome. Each panel compares WT Dcr2 with no Dcr2, an eGFP expressing Dcr2 negative treatment, or PAZ domain mutant M1-M4. The linear regression line is depicted within each plot, with the *R^2^* value indicating the fit of the model. **(B)** Heatmap displaying pairwise *R^2^* values from the linear regressions, comparing the similarity of differential coverage profile between treatments, as indicated. **(C)** Multidimensional scaling (MDS) plots representing the relationships between different size classes of vsiRNA reads from WT Dcr2, eGFP control and PAZ domain mutant Dcr2 M1-M4. Each plot corresponds to a specific size class of virus-derived small RNA: 20 nt, 21 nt, 22 nt, and 23 nt, as well as a combined plot of 20-23 nt. The axes represent the leading two log-fold change (logFC) dimensions and the percentage of variation indicated.

In addition to analyzing positional cleavage patterns of 21 nt vsiRNAs (Fig. 6A-B), we examined population-level patterns and diversity of SFV-derived small RNA populations for individual lengths of 20 nt, 21 nt, 22 nt, and 23 nt, as well as combined lengths of 20-23 nt by conducting a differential transcript abundance analysis using edgeR (Fig. 6C). Through the generation of a non-redundant list of unique small RNAs, we generated count tables and normalized for library size using the weighted trimmed mean of M-values (TMM) method. Through multi-dimensional scaling (MDS), plots we were able to visualize the relationships between the treatments. MDS plots are a form of dimensionality reduction; visually, the distances between points represent the dissimilarities or differences between the treatments. Specifically, these distances correspond to the leading log-fold changes between the treatments, which reflect the relative expression levels of vsiRNAs across conditions. The MDS plots revealed varying degrees of similarity between the WT and PAZ domain mutant Dcr2 treatments across different virus-derived small RNA size classes. Notably, the 21 nt vsiRNAs displayed distinct clustering patterns, indicating that specific mutations in the Dcr2 PAZ domain contribute to the altered distribution and abundance of these vsiRNA species.

## DISCUSSION

While the importance of the helicase domain has been demonstrated for both Dm and Aaeg Dcr2 antiviral responses (Gestuveo et al. 2022; Marques et al. 2013; Donelick et al. 2020), the relevance of the PAZ domain had not yet been investigated. Here, we identified conserved amino acids in Aaeg Dcr2 that inactivate PAZ domain function and antiviral activity. Critically, we observed that while inactivation did affect the production of 21 nt vsiRNA, it did not affect virus-derived small RNA production in general but broadened the length of the produced virus-derived small RNAs, similar to previous mutational analysis of Dm Dcr2 PAZ domain (Kandasamy and Fukunaga 2016). Thus, key observations hold true for viral dsRNAs and Aaeg Dcr2. The specific functions that are affected by the mutations we set in Aaeg Dcr2 PAZ will require further investigation. However, structural modelling (S1 Fig) suggests positioning mostly in the core of the PAZ domain and thus a role in the binding mechanism of the dsRNA termini. This is further confirmed by modelling Aaeg PAZ domain interactions with dsRNA (Supplemental Fig. S4). Suggesting that the location of dsRNA is not affected but most likely mutations destabilise binding and thus allow flexibility in the interaction, which in turn leads to variable small RNA lengths following cleavage. Importanly, the critical PAZ domain amino acids we described here in Aaeg Dcr2 are different from those previously identified in Dm Dcr2 (Kandasamy and Fukunaga 2016). Based on our and previous data that demonstrate the importance of the Dcr2 helicase domain in the exo-siRNA pathway, we propose that the helicase binds usually blunt-ended, long viral dsRNA and cleavage of these termini -which is likely to generate small RNAs of variable length-generates the first dsRNA terminus with 2 nt overhang and 5’ monophosphate that will be used by the PAZ domain to anchor the dsRNA strand and allow Dcr2 to produce mostly, though not exclusively, 21 nt vsiRNAs, and phase variation from occasional unprecise binding that allows the production of a diverse pool of vsiRNAs from dsRNA, may therefore start from early cleavage events onwards. Importantly, the size of 21 nt is critical and the most efficient siRNA length for silencing to target RNAs in the fly system (Kandasamy and Fukunaga 2016; Elbashir et al. 2001). Our data demonstrate that 21 nt vsiRNA are also optimal, and required for efficient silencing activity against an alphavirus infection in mosquito cells. While this has been assumed, here we provide experimental evidence that this is indeed the case.

In summary, this study demonstrates the critical role of the Aaeg Dcr2 PAZ domain in the production of specific 21 nt vsiRNA, but not small RNA production overall. Its critical contribution is in accurate vsiRNA length determination, which in turn is critical for the *Ae. aegypti* exo-siRNA to mount an antiviral response against SFV. Moreover, future work will determine how arboviruses of different families are targeted by Dcr2 and give us further insights into the nature of the viral dsRNA substrate.

## MATERIALS AND METHODS

### Cells

The Dcr2 knockout cell line Aag2-AF319 (subsequently abbreviated to AF319) used here is derived from *Ae. aegypti* Aag2-AF5 cells, a clone derived from the Aag2 cell line (provided by Kevin Maringer, The Pirbright Institute, UK) (Fredericks et al. 2019; Varjak et al. 2017). As previously described AF319 cells were cultured in Leibovitz’s L-15 medium with GlutaMax (Gibco) with 10% tryptose phosphate broth (TPB; Gibco), 10% fetal bovine serum (FBS; Gibco), and penicillin-streptomycin (pen-strep; 100 U/mL-100 µg/mL; Gibco) at 28 °C (Fredericks et al. 2019; Varjak et al. 2017). Cell lines are available as Aag2-AF319 (ECACC 19022602) and Aag2-AF5 (ECACC 19022601) through Public Health England. Baby hamster kidney (BHK-21) cells are a commonly used cell line available at the MRC-University of Glasgow Centre for Virus Research; these cells were grown in Glasgow Minimum Essential Medium (GMEM; Gibco) supplemented with 10% TPB, 10% newborn calf serum (Gibco), and pen-strep at 37 °C with 5% CO_2_.

### Viruses

Virus production from icDNA and plaque assay titrations were performed as previously described (Ulper et al. 2008; Varjak et al. 2017) to produce SFV4 (abbreviated to SFV; GenBank ID: KP699763) and SFV4(3H)-*FFLuc* (abbreviated to SFV-FFLuc; reporter virus expressing FFLuc).

### Plasmid generation

Mutations in the PAZ domain of Aaeg Dcr2 (GenBank ID: AAW48725) were created by In-Fusion cloning of synthetically generated PAZ domain mutant fragments into the previously generated pPUb-myc-Dcr2 (Varjak et al. 2017) following the manufacturer’s guidelines. Briefly, pPUb-myc-Dcr2 was linearised by PCR, removing the WT PAZ domain. Mutant PAZ domain fragments were PCR amplified using primers with extensions that were homologous to the linearised vector. In-Fusion cloning was then performed to create pPUb-myc-Dcr2 constructs containing the various mutants. Mutations were based on multiple sequence alignment and identification of conserved amino acids in *Ae. aegypti*, *D. melanogaster* and *Ae. albopictus*.

### Virus infections

For investigations of antiviral activity against SFV, AF319 cells were seeded at 2 x 10^5^ cells/well in 24-well plates and transfected with 1 µg pPUb plasmid expressing WT or mutant Dcr2 or eGFP (as control) after 24 h using Dharmafect2 (Horizon Discovery) following the recommended protocol. At 24 hpt, the cells were infected with SFV-FFLuc (MOI=0.1) and lysed at 48 hpi using 1X Passive Lysis Buffer (PLB; Promega). FFLuc levels were measured using a Luciferase Assay System (Promega) according to the manufacturer’s protocol in GloMax Multi-Detection System with Dual Injectors (Promega). For the production of samples for small RNA sequencing, 2.5 x 10^5^ cells/well AF319 cells were seeded in 8 wells of a 24 well plate, transfected as above after 24 h and infected with SFV (MOI=1) at 24 hpt for 48 h. For RNA extraction and follow up small RNA sequencing, cells from 7 wells were combined and lysed using TRIzol (Thermo Fisher Scientific) according to the manufacturer’s guidelines, with glycogen as a carrier. The remaining well was lysed in LDS protein lysis buffer and used for confirmatory western blot analysis.

### dsRNA production

FFLuc (dsFFLuc) and eGFP (dseGFP) were amplified using T7 RNA polymerase promoter-flanked primers (Varjak et al. 2017; McFarlane et al. 2020) and dsRNAs produced with MEGAscript RNAi kits (Thermo Fisher Scientific) by *in vitro* transcription as per manufacturer’s guidelines. Following *in vitr*o transcription, products were treated with DNaseI, and RNase A. dsRNA was purified by column purification.

### RNAi reporter assay

RNAi reporter assays were performed as previously described (Varjak et al. 2017) to assess the silencing activities of mutant Dcr2 compared to WT Dcr2. For this, 2 x 10^5^ AF319 cells/well were seeded in 24-well plates. After 24 h cells were co-transfected with 1 µg pPUb-based expression plasmid, 50 ng pGL3-PUb (Anderson et al. 2010) and 20 ng pPUb-RLuc (Alexander et al. 2023) together with either 20 ng dsRNA targeting FFLuc (dsFFLuc) or 20 ng dsRNA control (dseGFP) and lysed at 24 hpt in 1X PLB (Promega). To determine FFLuc/RLuc levels, dual luciferase assays were performed using a Dual-Luciferase Reporter Assay System (Promega) according to the manufacturer’s protocol in a GloMax Multi-Dectection System with Dual Injectors (Promega).

### Western blot analysis

Cell lysis was performed in 1X Bolt sample reducing agent and 1X Bolt LDS sample buffer (Invitrogen). The lysate was separated using a 4-12% Bis-Tris Plus gel (Invitrogen). For transfer onto a nitrocellulose membrane, a Trans-Blot SD Semi-Dry Transfer Cell (BioRad) was used. Membranes were blocked for a minimum of 1 h using 5% (w/v) non-fat dry milk powder in PBS-Tween (PBS with 0.05% Tween 20, PBS-T) and then washed three times for 10 min using PBS-T. As primary antibodies, mouse anti-myc tag antibody (1:2000, Abcam) and mouse anti-α tubulin antibody (1:2000; Sigma-Aldrich) in 5% (w/v) non-fat dry milk powder in PBS-T were used. After overnight incubation at 4 °C, membranes were again washed three times in PBS-T. This was followed by incubation of 1 h with goat anti-mouse DyeLight 800 (1:5000; Invitrogen) secondary antibody conjugated with a near fluorescent dye in 5% (w/v) non-fat dry milk powder in PBS-T. The membranes were again washed three times with PBS-T and a final wash with distilled water and analysed using an Odyssey CLx and Image Studio Lite v.5.2.5 (LI-COR Biosciences).

### Small RNA sequencing and data analysis

To investigate small RNA production, at least 1 µg isolated total RNA per condition were sent for analysis. Small RNA sequencing was performed at BGI-Tech Solutions (DNBSeq, UMI small library, SE50). with a minimum of 10 million clean reads, with final libraries resulting between 11-18 million reads (Supplemental Table S2).

### Small RNA analysis

Using fastp (v0.23.2) (Chen et al. 2018), adapters and low-quality reads from basecalled fastq files were trimmed, retaining reads between 18-32 nt. The distribution of read sizes and first position bias was calculated using the script ‘1_fastq_histogram.sh’ from the GitHub repository (rhparry/viral_sRNA_tools). Clean, trimmed reads were then mapped to the SFV genome (GenBank ID: KP699763) with Bowtie2 (v2.4.5) (Langmead and Salzberg 2012), using the sensitive mapping flag (—sensitive) as specified in the script ‘2_mapping_vrnas.sh’. Histograms of mapped read lengths and first base pair bias were generated using samtools (v1.16.1) and ‘3_bam_sRNA_histogram.sh’. Output BAM files were filtered to include only the 21 nt reads. Coverage statistics across the SFV genome were calculated using ‘4_viral_srna_coverage.sh’, which employs the bedtools genome coverage tool (v.2.27.1) (Quinlan and Hall 2010). For calculating the harmony metric, coverage at the 5’ end (−5 flag) of each read was normalized over the whole genome. Normalised coverage files were processed with awk ‘{f[NR]=$2;r[NR]=$3;sumf+=$2;sumr+=$3} END {for (i=1;i<=NR;i++) {printf “;%d %.6e\n”, i, (f[i]/sumf)-(r[i]/sumr)}}’ to calculate *d_i_* values for each position *i* of the SFV genome. Linear regression analyses for downstream analysis scatter plots of per position *d_i_* values of the SFV genome were analysed in GraphPad Prism (v10.0.2), and R^2^ values for all pairwise treatments were computed and presented as a heat map. Overlapping virus-derived small RNA pairs and overlap probabilities (z-score) were calculated iteratively from BAM files using the ‘signature.py’ small RNA signatures Python script, both single read lengths from 18-30 nt were calculated as per the following conditions (--minquery 18 --maxquery 18 --mintarget 18 --maxtarget 18 --minscope 1 --maxscope 18), collective 18-30 nt read lengths were computed as follows (--minquery 18 --maxquery 30 --mintarget 18 --maxtarget 30 -- minscope 1 --maxscope 30) (Antoniewski 2014). To examine the diversity and populations of SFV-derived small RNA species, unique 20-23 nt SFV mapped reads were treated as discrete virus-derived small RNAs. Counts tables for all extracted SFV-derived small RNAs from extracted fasta files from individual libraries were converted, processed, and enumerated in bash (v.3.2). The dimensions of count tables for 20-23 nt virus-derived small RNA species (n = 75710), 20 nt (n = 13430), 21 nt (n = 27592), 22 nt (n = 20536) and 23 nt (n = 14155) species. Differential gene expression analysis was performed using package edgeR (v.3.30.3) (Robinson et al. 2009) in R Studio (v.1.3.1073), with library size normalization using the weighted TMM method in calcNormFactors() and filtered for a minimum count of 10 with filterByExpr(). Relationships between samples were explored using an MDS plot of normalized count tables with plotMDS(top = 500) of all SFV-derived small RNA species.

### Protein structure predictions

The WT sequence of Aaeg Dcr2 (GenBank ID: AAW48725) was used as a query for structure prediction. AlphaFold 2 algorithm (Jumper et al. 2021) was run in monomer mode using a local install of version 2.3.2 without any restrictions. The resulting models were examined and, in accordance with overall model quality predictions, summarised in mean pLDDT value with the highest quality model selected for further analysis. Structural analyses were performed in PyMOL Molecular Graphics System (version 4.5.0, Schrödinger, LLC). Domain positions were assigned and labelled by color based on GenBank sequence annotations, and residues mutated were highlighted.

*Ae. aegypti* Dcr2-dsRNA interactions were assessed using AlphaFold 3. The WT sequence of *Ae. aegypti* Dcr2 (GenBank ID: AAW48725) and derived mutant sequences (M1-M4) were modelled as interacting with dsRNA. The 53 bp dsRNA sequence was obtained from PDB ID: 7W0E for *D. melanogaster* Dcr2 in the active dicing state (Su et al. 2022). Sequences for WT or M1-M4 *Ae. aegypti* Dcr2 and each mutant were fed into AlphaFold 3 webserver (Abramson et al. 2024) along with the dsRNA sequence to predict protein structures and their interaction with dsRNA. Molecular graphics and analysis of structures were performed using UCSF ChimeraX (Pettersen et al. 2021).

### Data analyses

Virus-derived small RNA metrics, coverage and overlapping pairs analysis were visualized using GraphPad Prism (v10.0.2).

## DATA DEPOSITION

Data underlying figures are available under http://dx.doi.org/10.5525/gla.researchdata.1479. Small RNA sequencing data generated for this study have been deposited in the NCBI Sequence Read Archive (SRA), available under accession number PRJNA1014654. Scripts utilized for small RNA analysis are available from the viral_sRNA_tools GitHub repository at: https://github.com/rhparry/viral_sRNA_tools

## SUPPLEMENTAL MATERIAL

Supplemental material is available for this article.

## ACKNOWLEDGMENTS

This study was supported by the UK Medical Research Council (MC_UU_12014/8, MC_UU_00034/4) (A.K.) and (MR/R021562/1, MC_UU_00034/2) (A.C.); European Research Council (ERC) Consolidator Grant ‘vRNP-capture’ N# 101001634 (A.C.); German Centre for Infection Research (DZIF) (TTU 01.708) (E.S.); DFG (project 497659464) (M.R.); Wellcome Trust/Royal Society Sir Henry Dale Fellowship (210462/Z/18/Z) (B.B.); Overseas Scholarship Scheme for PhD by Higher Education Commission of Pakistan (R.A.).

Author contributions are recognised as follows. Conceptualization: M.R., R.P., M.M., R.J.G., A. C., M.V., L.R., E.S., A.K.; Methodology: M.M., M.R., R.P., R.A., R.J.G., L.R.; Formal analysis: M.R., R.P., M.M., R.A., L.R.; Investigation: M.R., R.P., M.M., R. A., L.R.; Validation: M.R., R.P., M.M., R.A., L.R..; Resources: A.A.K., B.B., A.C., L.R., E.S., A.K.; Writing - Original Draft Preparation: M.R., R.P., M.M., R.J.G., R.A., A.C., L.R., E.S., A.K.; Writing - Review and Editing: M.R., R.P., M.M., R.J.G., R.A., A.A.K., B.B., M.V, A.C., L.R., E.S., A.K..; Visualization: M.R., R.P, M.M., R.A., L.R..; Data Curation: M.R., R.P., M.M., E.S., A.K.; Supervision: M.M., A.A.K., B.B., A.C., E.S., A.K.; Project Administration: M.M., E.S, A.K.; Funding Acquisition: R.A., B.B., A.C., E.S., A.K.

